# Peptidylarginine deiminase IV (PADI4) is not essential for cell-autonomous HSC maintenance and normal haematopoiesis

**DOI:** 10.1101/2021.04.13.439513

**Authors:** Christine Young, John R. Russell, Hannah Lawson, Christopher Mapperley, Kamil R. Kranc, Maria A. Christophorou

## Abstract

Peptidylarginine deiminases (PADIs, or PADs) are emerging as key regulators of human physiology and pathophysiology. The nuclear deiminase PADI4 regulates embryonic stem cell pluripotency, however its role in adult stem cells is unknown. PADI4 is expressed most highly in the bone marrow (BM), where it is found as part of a self-renewal-associated gene signature and shown to modulate the function of critical transcriptional regulators such as Tal1 and c-Myc, suggesting that it regulates haematopoietic development or regeneration. We investigated the functional significance of PADI4 in haematopoietic stem cell (HSC) biology and normal haematopoiesis. We employed two conditional mouse models of tissue-specific *Padi4* ablation, where *Padi4* was completely deleted either after the emergence of HSCs, or acutely in the BM of adult mice. We found that loss of PADI4 does not significantly affect HSC self-renewal or differentiation potential upon injury or serial transplantation, nor does it lead to exhaustion or premature ageing of HSCs. Thus, surprisingly, PADI4 is dispensable for cell-autonomous HSC maintenance, differentiation and haematopoietic regeneration. This work has important implications for the clinical use of PADI4 inhibitors as therapeutic agents in autoimmunity and cancer.

**Key Points:**

- PADI4 is dispensable for steady-state and post-transplantation haematopoiesis
- HSCs do not require intrinsic PADI4 activity to respond to haematopoietic injury
- PADI4 deficiency does not lead to premature HSC ageing or exhaustion

## Introduction

Haematopoietic stem cells (HSCs) possess self-renewal capacity and multi-lineage differentiation potential, and are therefore able to replenish all blood cell lineages, sustaining normal and post-injury haematopoiesis. In addition to transcription factors, which directly facilitate or inhibit gene transcription, one central mechanism involved in stem cell fate decisions is the modulation of expression of stem cell and differentiation genes achieved *via* epigenetic mechanisms such as histone modifications. Indeed, several studies have shown that histone-modifying enzymes are essential for normal haematopoiesis^1,2,3,4,5,6,7^.

Peptidylarginine deiminase (PADI) enzymes catalyze citrullination, the post-translational conversion of arginine residues within a protein to the non-coded amino acid citrulline. The five PADI family members are structurally similar and likely to operate *via* common regulatory mechanisms^8,9^, but they show varying tissue distributions and sub-cellular localisations, suggesting that they have specific organismal roles. They have been shown to regulate fundamental molecular and cellular processes such as gene expression, chromatin compaction, nerve myelination, the innate immune response and pluripotency^10,11,12,13,14,15,16,17^, while lack of PADI activity leads to infertility and defects in neurodevelopment and embryo development^11,18,19^. PADI4 citrullinates core and linker histone proteins and has well-established roles in the regulation of gene transcription and chromatin compaction. We previously showed that PADI4 mediates the establishment of pluripotency^11^, however it is unknown whether it has a role in adult stem cells. Out of all mammalian tissues, *Padi4* is most highly expressed in the bone marrow (BM) and is one of the top 50 genes associated with self-renewal, as determined by the fact that it is down-regulated upon differentiation to all multi-lineage progenitors, but up-regulated again in leukaemia stem cells^20^. More recent studies showed that haematopoietic multipotent progenitor cells from *Padi4*-null mice exhibit increased proliferation and that PADI4 is a co-activator of Tal1, a critical transcriptional regulator in the haematopoietic system^21,22^. Taken together, these findings suggest that PADI4 functions in the regulation of haematopoiesis.

To understand the role of PADI4 in normal haematopoiesis, HSC maintenance, haematopoietic regeneration and ageing, we constitutively and inducibly deleted *Padi4* from the haematopoietic system. We demonstrate that HSCs do not require PADI4 to self-renew, sustain long-term multilineage haematopoiesis or respond to haematopoietic injury. Moreover, by investigating long-term consequences of *Padi4* deletion, we show that *Padi4* loss does not lead to HSC exhaustion or premature ageing.

## Results and Discussion

To determine the functional significance of PADI4 in steady-state haematopoiesis and HSC self-renewal, we deleted *Padi4* specifically from the haematopoietic system using the *Vav-iCre* system. *Vav-iCre* mice^23^ constitutively express the codon-improved Cre (iCre)^24^ driven by the *Vav* regulatory elements^25^, resulting in haematopoietic-specific gene deletion shortly after the emergence of definitive HSCs^26^ and ensuring recombination in all HSCs^27,28^. We bred these mice to *Padi4*^*fl/fl*^ mice^29^, in which *Padi4* exons 9 and 10 are flanked by *loxP* sites. These exons contain aspartate 352, which is part of the active site, as well as four additional residues (Q351, E353, E355, D371), which are essential for Ca^2+^ binding and activation of the enzyme^8^. The resulting *Padi4*^*fl/fl*^*;Vav-iCre* mice (referred to as *Padi4*^CKO^ hereafter) completely lack PADI4 protein expression in the BM (Figure **1A**). These mice were compared to *Padi4*^fl/fl^ mice (referred to as *Padi4*^CTL^) in all subsequent analyses. *Padi4*^CKO^ and *Padi4*^CTL^ mice showed normal Mendelian distribution, had comparable survival and did not display any obvious defects. To enumerate cells at different levels of the haematopoietic differentiation hierarchy, we next carried out immunophenotypic analyses of *Padi4*^CKO^ mice. In agreement with a previous report^21^, *Padi4*^CKO^ mice had increased numbers of Lin^-^Sca-1^+^c-Kit^+^ (LSK) cells but similar numbers of total white blood cells (WBC) and lineage restricted myeloid and erythroid of Lin^-^Sca-1^-^c-Kit^+^ (LK) progenitor cells compared to *Padi4*^CTL^ mice (Figure **1B**). Further analysis of the LSK compartment showed normal numbers of LSKCD48^-^CD150^+^ HSCs, LSKCD48^-^CD150^-^ multipotent progenitors (MPPs), LKSCD48^+^CD150^+^ primitive haematopoietic progenitors (HPC-2) and an increase in LSKCD48^+^CD150^-^ haematopoietic progenitor cell-1 (HPC-1) population in *Padi4*^CKO^ mice compared to *Padi4*^CTL^ mice (Figure **1C**). We observed an increase in common lymphoid progenitors (Lin^-^c-Kit^lo^Sca^lo^IL7Rα^+^cells) in *Padi4*^CKO^ mice compared to *Padi4*^CTL^ mice (Figure **1D**). However, the numbers of myeloid and erythroid progenitors, CD11b^+^Gr1^-^ and CD11b^+^Gr1^+^ differentiated myeloid cells, CD19^+^ B cells and Ter119^+^ erythroid cells, were comparable in the BM between *Padi4*^CKO^ and *Padi4*^CTL^ mice (Figure **1E**). These results were mirrored in *in vitro* colony forming cell (CFC) assays where there was no difference in colony counts between the two genotypes (Figure **1F**). In addition, numbers of thymic T-cells were unaffected (Figure **1G**). Analysis of peripheral blood (PB) showed that the numbers of circulating blood cells and haemoglobin parameters were completely unaffected by *Padi4* deletion (Supplementary Figure 1A). Analysis of the spleens showed a modest increase in differentiated cells in *Padi4*^CKO^ mice with an overall increase in WBC counts, indicating extramedullary haematopoiesis (Figure **1H**). In conclusion, despite mild extramedullary haematopoiesis, *Padi4* deletion has no major impact on BM steady-state haematopoiesis.

**Figure 1.**
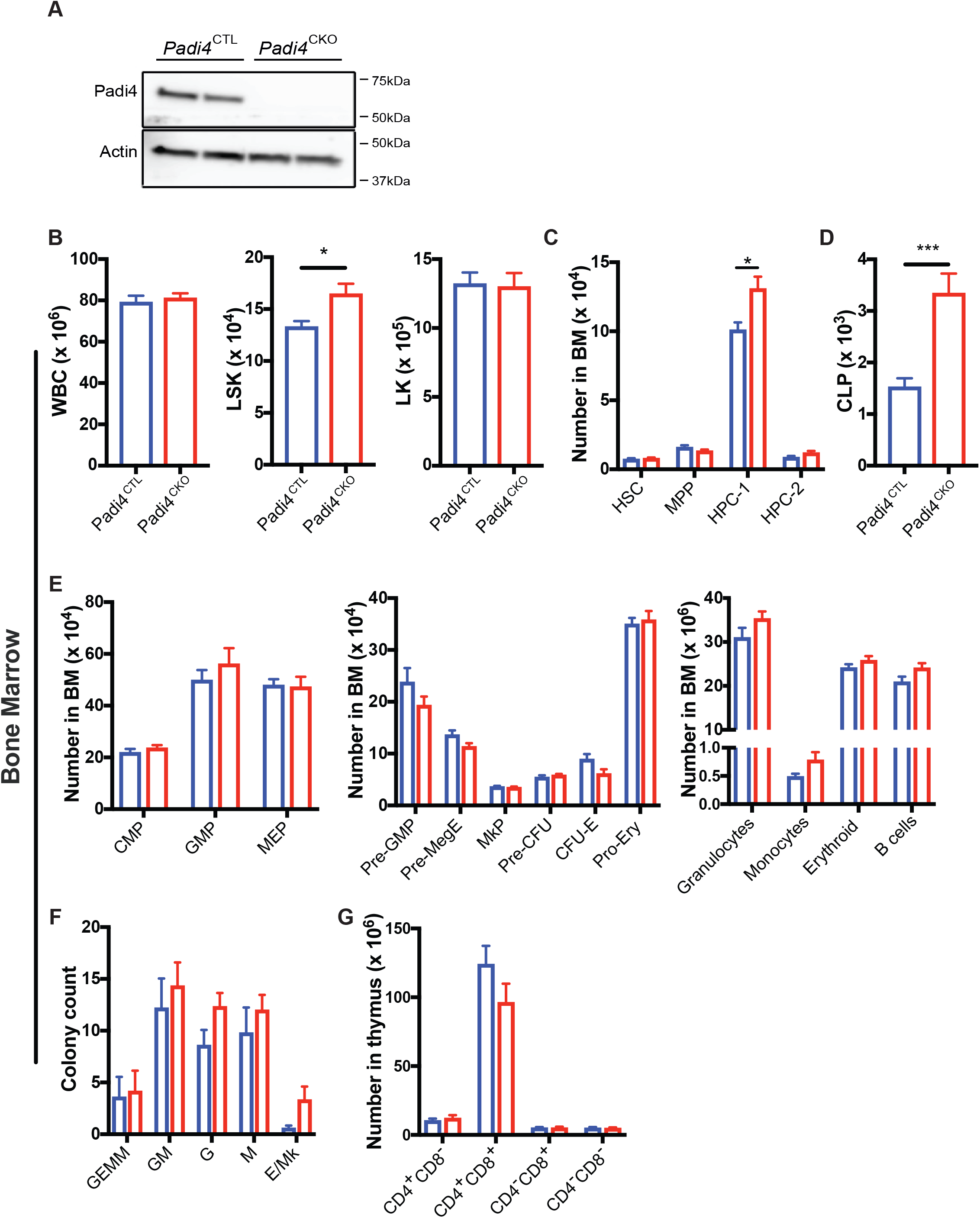
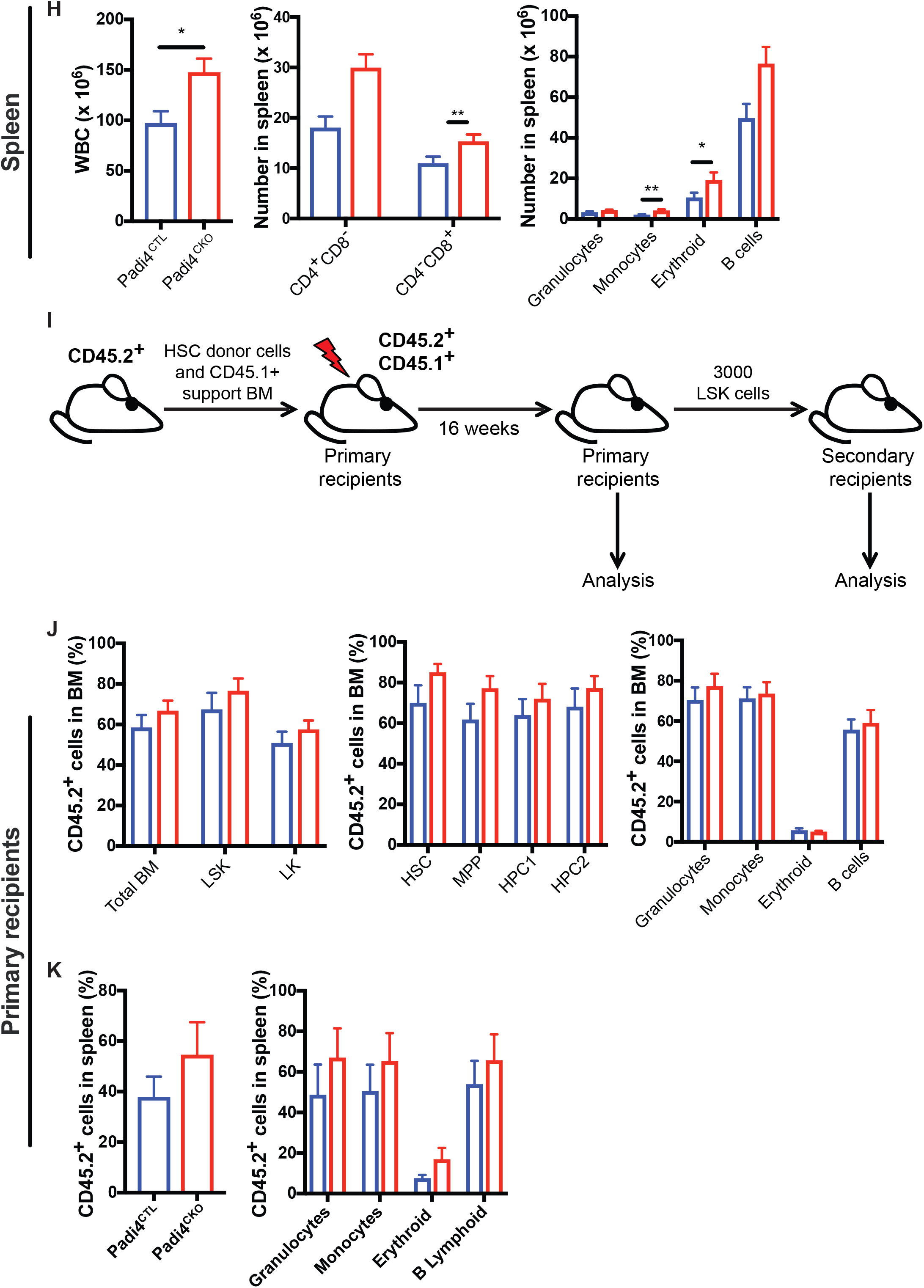
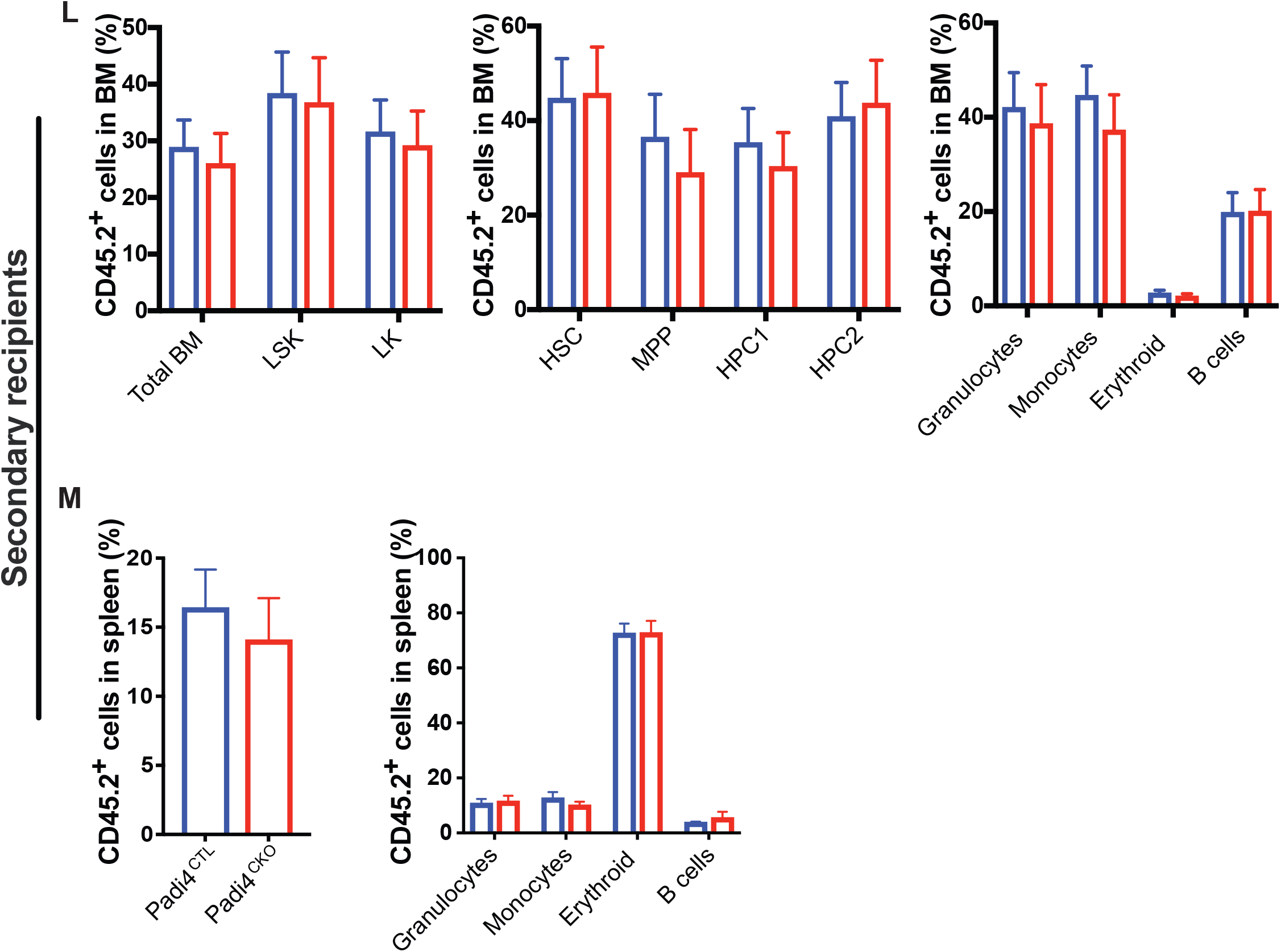
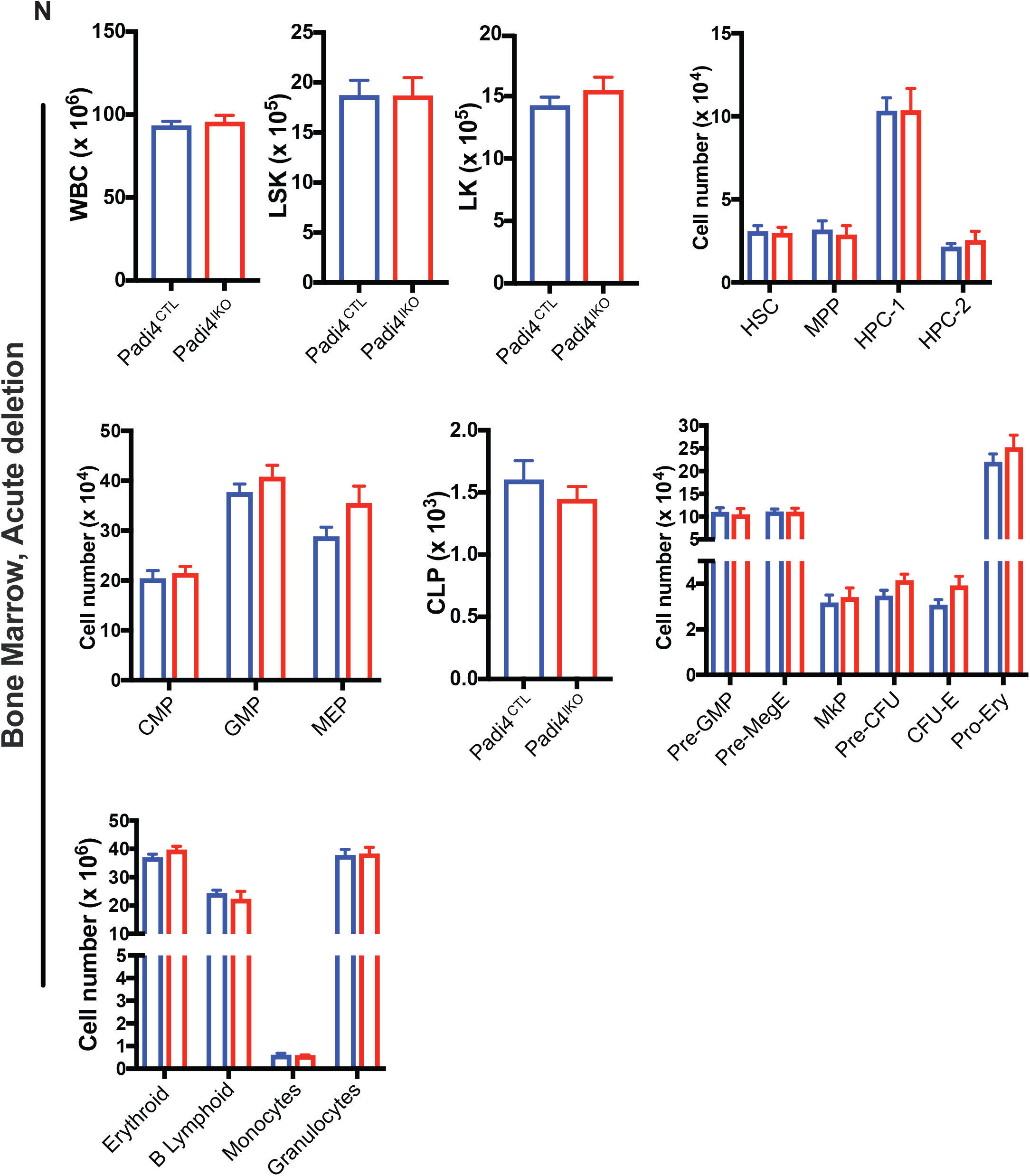
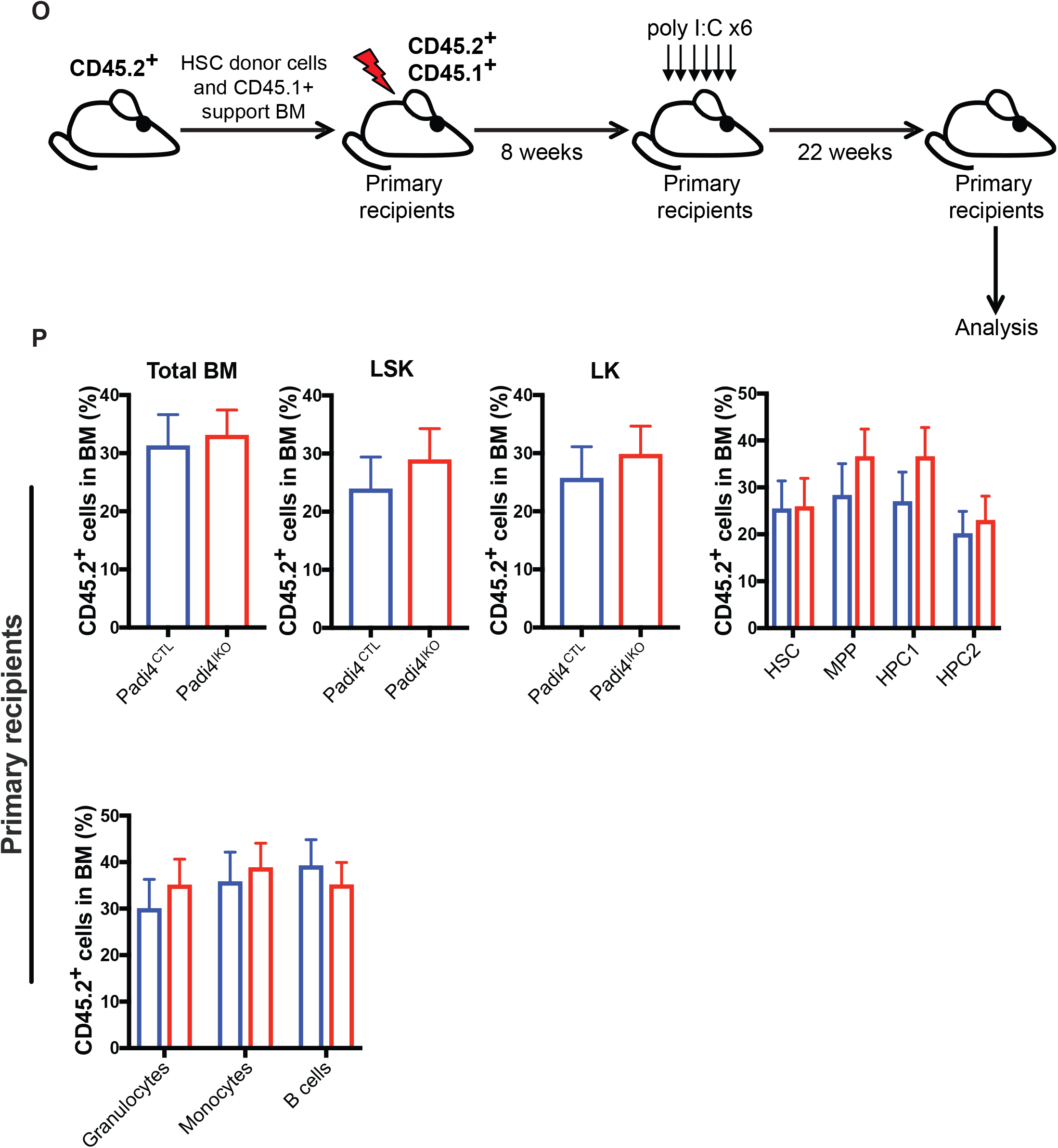
Haematopoiesis-specific deletion of PADI4 has no major impact on steady-state haematopoiesis or BM reconstitution potential following serial transplantations. **(A)** Immunoblot analysis of mouse PADI4 in total BM extracts from *Padi4*^CTL^ and *Padi4*^CKO^ mice. Actin presented as a loading control. **(B-G)** Immunophenotypic analysis of bone marrow from 8-12 week old mice; total number of **(B)** WBC, LSK, and LK cells **(C)** HSC, MPP, HPC-1 and HPC-2 cells. *Padi4*^CTL^, n = 9; *Padi4*^CKO^, n = 9. **(D**,**E)** Total number of lymphoid, myeloid and erythroid progenitor cells. **(D)** CLP, **(E)** CMP, GMP, MEP, Pre-GMP, Pre-MegE, MkP, Pre-CFU, CFU-E, Pro-Ery and differentiated cell populations (Granulocytes, Monocytes, Erythroid and B cells). *Padi4*^CTL^, n = 9; *Padi4*^CKO^, n = 9. **(F)** CFC assay with BM cells. *Padi4*^CTL^, n = 5; *Padi4*^CKO^, n = 6. **(G)** Total number of thymic T cells. *Padi4*^CTL^, n = 9; *Padi4*^CKO^, n = 9. **(H)** Immunophenotypic analysis of spleen from 8-12 week old mice; total number of WBC, T cells and differentiated cell populations (Granulocytes, Monocytes, Erythroid and B cells). *Padi4*^CTL^, n = 9; *Padi4*^CKO^, n = 9. **(I)** Experimental design for BM transplantation experiments. 200 CD45.2+ BM HSCs from C57BL/6 *Padi4*^CTL^ or *Padi4*^CKO^ mice were transplanted into primary recipient mice and monitored for 16 weeks. Following this, a cohort of mice were sacrificed for analysis at 16 weeks post transplantation and bone marrow was transplanted to secondary recipients. **(J)** Percentage of donor-derived CD45.2+ cells in total BM, LSK, LK, HSC, MPP, HPC-1, HPC-2 and differentiated cell populations (Granulocytes, Monocytes, Erythroid and B cells). *Padi4*^CTL^, n = 14; *Padi4*^CKO^, n = 13. **(K)** Contribution of donor-derived CD45.2+ cell population to total spleen WBC count and differentiated cell populations of primary recipients. *Padi4*^CTL^, n = 6; *Padi4*^CKO^, n = 6. **(L-M)** Secondary recipient mice were transplanted with 3000 sorted CD45.2+ BM LSK cells from primary recipients sacrificed at 16 weeks. **(L)** Percentage of donor-derived CD45.2+ cells in total BM, LSK, LK, HSC, MPP, HPC-1, HPC-1 and differentiated cell lineages (Granulocytes, Monocytes, Erythroid and B cells). *Padi4*^CTL^, n = 19; *Padi4*^CKO^, n = 19. 2-4 donors were used per genotype. **(M)** Contribution of donor-derived CD45.2+ cell population to spleen WBC and differentiated cells of secondary recipients. *Padi4*^CTL^, n = 19; *Padi4*^CKO^, n = 19. **(N-P)**. Acute deletion of *Padi4* in adult HSCs. *Padi4*^CTL^ and *Padi4*^IKO^ mice received 6 x IP injection of Poly I:C to induce deletion of Padi4. **(N)** Immunophenotypic analysis performed 4 weeks following the final injection. Total number of: WBC, LSK, LK, HSC, MPP, HPC-1 and HPC-2 cells; myeloid, erythroid and lymphoid progenitor cells: GMP, MEP, CMP, CLP Pre-GMP, Pre-MegE, MkP, Pre-CFU, CFU-E, Pro-Ery; differentiated granulocytes, monocytes, B cells and erythroid cells in the BM. n = 5 -8 per genotype. **(O)** Schematic of experimental procedure for transplantation of *Padi4*^CTL^ and *Padi4*^IKO^ BM cells. 2 × 10^5^ unfractionated CD45.2+ BM cells from untreated *Padi4*^CTL^ and *Padi4*^IKO^ C57BL/6 (8– 12 wk old) mice were mixed with 2 × 10^5^ CD45.1+ WT BM cells and transplanted into lethally irradiated CD45.1+/CD45.2+ recipients. 8 wk after transplantation, the recipients received six doses of Poly I:C. **(P)** Percentage of donor-derived CD45.2+ cells in the BM of recipient mice: LSK, LK, HSC, MPP, HPC-1, HPC-1 and differentiated cell lineages (Granulocytes, Monocytes and B cells). n = 15–21 recipients per genotype. n = 3-4 donors per genotype. All data are mean ± SEM. *, P < 0.05; **, P < 0.01; ***, P < 0.001, ****, P < 0.0001 (Mann-Whitney U test).

To assess the requirement for *Padi4* in HSC maintenance, we performed competitive HSC transplantation assays. CD45.2^+^LSKCD48^-^CD150^+^ HSCs sorted from *Padi4*^CKO^ and control (*Padi4*^CTL^) mice were competitively transplanted into lethally irradiated wild-type syngeneic CD45.1^+^/CD45.2^+^ recipients (Figure **1I**). Peripheral blood analysis showed no difference in CD45.2^+^ donor-derived chimerism in primary recipients of the *Padi4*^CKO^ HSCs when compared to recipients of *Padi4*^CTL^ HSCs (Supplementary Figure 2A). BM analysis at 16 weeks post-transplant showed that HSCs of both genotypes efficiently reconstituted long-term multilineage haematopoiesis, while donor-derived cells contributed equally to BM HSC and primitive cell compartments of the primary recipient mice (Figure **1J**). No difference in CD45.2^+^ cell engraftment was found in the spleen of recipient mice and both *Padi4*^CTL^ and *Padi4*^CKO^ donor derived cells contributed equally to differentiated cell populations in the spleen (Figure **1K**). Moreover, *Padi4*^CKO^ LSK cells sustained long-term BM reconstitution in secondary recipients comparably to *Padi4*^CTL^ LSK cells (Figure **1L**), while equal engraftment of CD45.2^+^ cell engraftment was observed in spleen (Figure **1L**) of the secondary recipients. In addition, no difference in CD45.2^+^ cell engraftment was found in PB (Supplementary Figure 2B). These experiments revealed that HSCs do not require *Padi4* to self-renew and sustain long-term multi-lineage haematopoiesis upon transplantation.

Given that *Vav-iCre* recombines in the embryo soon after the emergence of definitive HSCs, it is possible that that deletion of *Padi4* during haematopoietic development is compensated for by the activity of another member of the peptidylarginine deiminase family. Indeed, although PADI4 is the only predominantly nuclear member of the PADI family, PADI2 and PADI1 have also been shown to act in the nucleus^17,30,31^ and regulate gene expression. To rule out the possibility of compensation, we examined HSC maintenance following acute *Padi4* deletion in adult mice. We generated *Padi4*^*fl/fl*^*;Mx1-Cre* mice (referred to as *Padi4*^IKO^ hereafter) in which efficient recombination in the BM is induced by treatment with Poly I:C^32^. *Padi4*^IKO^ and *Padi4*^CTL^ mice received 6 injections of 300 µg Poly I:C, on every other day, and were culled and analysed 4 weeks following the final administration. We found that acute deletion of *Padi4* had no effect on any bone marrow cell compartment analysed (Figure **1N**). To test whether acute *Padi4* deletion affects post-transplantation haematopoietic reconstitution, we transplanted CD45.2^+^ unfractionated BM cells from *Padi4*^IKO^ or *Padi4*^CTL^ mice with support CD45.1^+^ BM cells to lethally irradiated recipient mice. Following efficient CD45.2^+^ cell engraftment (8 weeks post transplantation), the mice received 6 doses of Poly I:C resulting in efficient *Padi4* deletion in donor-derived CD45.2^+^ cells (Figure **1O** and Supplementary Figure 3A). The contribution of CD45.2^+^ cells to the PB of the recipients was quantified at 4, 8, 12, 18 and 22 weeks post transplantation and no differences were observed (Supplementary Figure 3B). The donor-derived contribution of CD45.2^+^ cells to BM cell compartments of the recipients, including total WBC, LSK, LK and HSCs of the recipients was similar regardless of the genotype of transplanted CD45.2^+^ cells, as was the CD45.2^+^ cell contribution to differentiated cell lineages in the BM (Figure **1P**). Therefore, compensation from another of the PADI family members can be excluded.

To test the role of *Padi4* in HSC regenerative capacity upon haematopoietic injury, we treated adult (8-12 week) *Padi4*^CKO^ and *Padi4*^CTL^ mice with 5-fluorouracil (5-FU). Mice received 3 injections of 5-FU 10 days apart and BM was analysed 10 days following the final dose (Figure **2A**). We observed no difference in numbers of HSCs, LSK, LK cell compartments or differentiated cells in the BM of *Padi4*^CKO^ mice compared to *Padi4*^CTL^ mice (Figure **2B** and Supplementary Figure 4). Therefore, PADI4 is dispensable for the ability of HSCs to respond to haematopoietic stress.

To investigate the long-term effects of Padi4 deletion in the haematopoietic system, we carried out immunophenotypic analyses of *Padi4*^CKO^ mice aged up to 1 year, as well as BM reconstitution experiments using HSCs from these aged mice (Figure **2C,D**). Analysis of BM and spleens of 1 year-old *Padi4*^CKO^ mice and *Padi4*^CTL^ mice showed no difference in cell counts of total WBC, primitive LSK compartments, LK progenitor populations or differentiated cell populations (Figure **2C**). In addition, *in vitro* CFC assays using aged BM from *Padi4*^CKO^ and *Padi4*^CTL^ mice also showed no difference in CFC colony count between the genotypes (Figure **2C**). In fact, the differences observed in young mice (Figure **1B,C**,**H**) were not observed upon ageing. To assess the self-renewal potential of aged *Padi4*^CKO^ HSCs, we transplanted sorted HSCs from 1-year old *Padi4*^CKO^ and *Padi4*^CTL^ mice into primary recipients and analysed the bone marrow of the recipients at 36 weeks post-transplantation. Mice transplanted with *Padi4*^CKO^ HSCs showed a small decrease in the contribution of donor derived CD45.2^+^ cells to the LK and HPC1 progenitor populations as well as a decrease in differentiated granulocytes (Figure **2D**). No differences in the ability of *Padi4*^CKO^ HSCs to contribute to spleen or peripheral blood cells was found (Figure **2D** and Supplementary Figure 5). Taken together, these results show that PADI4 is dispensable for long-term cell-autonomous HSC maintenance and normal haematopoiesis.

This study demonstrates that PADI4 is not required for steady-state haematopoiesis, long-term self-renewal of HSCs, efficient reconstitution of multi-lineage haematopoiesis in serial transplantation assays or response of HSCs to haematopoietic injury. This is surprising in the face of evidence that arginine methyltransferases, which catalyse protein arginine methylation, a modification that is antagonistic to citrullination^33,34^, have clear roles in the regulation of haematopoiesis^67^. PADI4 is expressed in haematopoietic and leukaemia stem cells, but not in committed haematopoietic progenitors^20^ and was shown to act as a co-activator of Tal1, a key transcriptional regulator in haematopoiesis, specifically by counteracting histone arginine methylation^22^. A previous study conducted using *Padi4*-null mice^21^ suggested that PADI4 regulates the proliferation of multipotent stem cells in the BM, as such mice showed increased numbers of LSK cells. Our experiments using haematopoiesis-specific deletion of *Padi4* replicate this phenotype but our comprehensive analyses of normal haematopoiesis and HSC maintenance under steady-state conditions and upon stress show that *Padi4* deletion does not affect any aspects of haematopoiesis, despite a transient increase in some of the multipotent progenitor cell populations. We therefore conclude that HSCs do not require intrinsic PADI4 activity for their cell-autonomous functions. As significant efforts are underway to generate potent and specific inhibitors of PADI4 as a therapeutic approach against autoimmune disorders and cancer^35^, this study indicates that these can be used systemically without adverse effects to the haematopoietic system.

## Materials and Methods

### Mice

All experiments on animals were performed under UK Home Office authorisation. All mice were of C57BL/6 genetic background. *Padi4*^fl/fl^ mice^29^ were a kind gift from the Mowen lab. *Vav-iCre*^23^, *Mx1-Cre*^36^, have been described previously^37^. All transgenic and knockout mice were CD45.2^+^. Congenic recipient mice were CD45.1^+^/CD45.2^+^. Sex-matched 8 to 12 week-old mice were used throughout.

### Flow cytometry

All BM cells were prepared and analyzed as described previously^37–41^. BM cells were isolated by crushing tibias and femurs using a pestle and mortar. Cell suspensions were passed through a 70µm strainer. PB was collected in EDTA coated microvettes. Spleen and thymus were homogenised and passed through a cell strainer. Single cell suspensions were incubated with Fc block and then stained with antibodies. For HSC cell analyses, unfractionated BM cells were stained with a lineage marker cocktail containing biotin-conjugated anti-CD4, anti-CD5, anti-CD8a, anti-CD11b, anti-B220, anti-Gr-1 and anti-Ter119 antibodies together with APC-conjugated anti-c-Kit, FITC-conjugated anti-Sca-1, PE-conjugated anti-CD48 and PE-Cy7-conjugated anti-CD150 antibodies. Biotin-conjugated antibodies were then stained with Pacific Blue-conjugated streptavidin. For pan-lineage progenitor cell staining, cells were stained with the lineage marker cocktail described above together with APC-conjugated anti-c-kit, PE-Cy7-conjugated anti-Sca-1, BV-421-conjugated anti-CD127, FITC-conjugated anti-CD34, PE-conjugated anti-CD135 and APC-Cy7-conjugated anti-CD16/32. For myeloid/T lymphoid restricted progenitors, cells were stained with a lineage marker cocktail containing biotin-conjugated anti-CD4, anti-CD5, anti-CD8a, anti-Mac-1, anti-B220, anti-CD19 and anti-Gr-1 together with BV-510-conjugated anti-c-kit, Pacific Blue-conjugated anti-Sca-1, PE-Cy7-conjugated anti-CD150, APC-Cy7-conjugated anti-CD16/32, APC-conjugated anti-CD41, PE-conjugated anti-CD105 and FITC-conjugated anti-Ter119. Biotin-conjugated antibodies were then stained with PerCP-conjugated streptavidin. For analyses of differentiated cells, spleen, BM or PB cell suspensions were stained with APC-Cy7-conjugated anti-CD19 antibody for B cells; Pacific Blue-conjugated anti-CD11b and PE-Cy7-conjugated anti-Gr-1 for myeloid cells; APC-conjugated anti-CD8 antibodies and PE-conjugated anti-CD4 antibodies for T cell analysis (spleen and PB); FITC-conjugated anti-Ter119 and PE-conjugated anti-CD71 (BM).

To distinguish CD45.2^+^-donor derived cells in the BM and spleen of transplanted mice, BV711-conjugated anti-CD45.1 and Pacific Blue-conjugated anti-CD45.2 antibodies were used. For HSC staining in transplanted mice, the remainder of the staining was as described above. For analyses of differentiated cells in BM and spleen of transplanted mice, myeloid cells were stained with PE-conjugated anti-CD11b, PE-Cy7-conjugated anti-Gr-1 and FITC-conjugated anti-Ter119 for erythroid cells. B Lymphoid cells were stained with APC-Cy7-conjugated anti-CD19. PB of transplanted mice was stained with FITC-conjugated anti-CD45.1, Pacific Blue-conjugated anti-CD45.2, PE-conjugated anti-CD4 and-CD8a, PE-Cy7-conjugated anti-Gr-1, APC-conjugated anti-CD11b, and APC-Cy7-conjugated anti-CD19.

Flow cytometry analyses were performed using a LSRFortessa (BD). Cell sorting was performed on a FACSAria Fusion (BD).

### Colony forming cells (CFC) assays

CFC assays were carried out using MethoCult™ M3434 (STEMCELL Technologies) methylcellulose medium. Two technical replicates were used per each biological replicate in each experiment. BM cells were plated for 10 days before colony types were identified and counted.

### Blood profiling

Blood was collected via cardiac puncture into an EDTA coated microvette and analysed on a Celltaq Haematology analyser (Nihon Kohden).

### Syngeneic transplantation assays

CD45.1^+^/CD45.2^+^ C57BL/6 recipient mice were lethally irradiated using a split dose of 11 Gy (two doses of 5.5 Gy administered at least 4 hours apart) at an average rate of 0.58 Gy/min using a Cesium 137 GammaCell 40 irradiator. For primary transplantations 200 LSKCD48^-^CD150^+^ HSCs (per recipient) sorted from BM of the donor mice were mixed with 200,000 unfractionated support CD45.1^+^ wild type BM cells and transferred into lethally irradiated CD45.1^+^/CD45.2^+^ recipients. For secondary transplantations 2,000-3,000 CD45.2^+^ LSK cells sorted from BM of primary recipients were mixed with 200,000 unfractionated support CD45.1^+^ wild-type BM cells and re-transplanted. For all except ageing experiments, primary and secondary recipients were culled and analysed 16-20 weeks post transplant.

### Poly I:C administration

Both *Padi4*^CTL^ and *Padi4*^IKO^ *(Padi4;Mx1-Cre)* transgenic and CD45.1^+^/CD45.2^+^ C57BL/6 recipient mice were injected intraperitoneally with poly I:C. *Padi4*^*CTL*^ *and Padi4*^IKO^ mice received 6 injections of 300 µg Poly I:C, on every other day. Mice were culled and analysed 1 month following the final administration. Recipient mice received one injection every other day with 300 µg Poly I:C (GE Healthcare) for a total of 6 doses starting 8 weeks after transplantation as previously described^38,40,41^.

### 5-FU administration

Both *Padi4*^*CTL*^ and *Padi4*^*CKO*^ transgenic mice were injected intraperitoneally with 5-FU. Mice were weighed on the day of administration and received 3 injections of 150 mg/kg 10 days apart. Mice were culled and analysed 10 days following the final administration.

### Western blotting

Proteins extracted from *Padi4*^*CTL*^ and *Padi4*^*CKO*^ were subjected to SDS–PAGE (4–20% Mini-PROTEAN^®^ TGX™ Precast gel, Biorad) and then transferred onto a nitrocellulose membrane. Membranes were blocked in 5% BSA-TBST (TBS with 0.1% Tween20) and probed with anti-Padi4 (Abcam ab214810, 1:1000, O/N at 4°C) and anti-Actin (Santa Cruz, sc-1616, 1:500, O/N at 4°C). After incubation with appropriate horseradish peroxidase-coupled secondary antibody, proteins were detected with SuperSignal™ West Pico PLUS Chemiluminescent Substrate (ThermoFisher Scientific) and acquired on the ImageQuant LAS (GE Healthcare Life Sciences).

### Genotyping

DNA was extracted from bone marrow taken from recipient mice transplanted with *Padi4*^CTL^, *Padi4*^CKO^ or *Padi4*^IKO^ mice. PCR was performed using primers specific for *Padi4* deletion. Forward primer: 5’-CAG GAG GTG TAC GTG TGC A-3’. Reverse primer: 5’-AGT CCA GCT GAC CCT GAA C-3’. Expected band sizes: Wild-type *Padi4* allele: 104bp; Floxed *Padi4* allele: 160bp; Knock-out *Padi4* allele: 215bp.

### Statistical analysis

Statistical significance was determined using Mann-Whitney or One-Way ANOVA on Graphpad V 8 software.

## Figure Legends

**Figure 2. Haematopoiesis-specific deletion of PADI4 has no major impact on BM reconstitution potential following haematopoietic injury and does not cause haematopoietic stem cell exhaustion in ageing mice. (A)** Experimental design of haematopoietic injury approach. *Padi4*^CTL^ and *Padi4*^CKO^ mice received a 3x 5-FU injections at 150 mg/kg 10 days apart and were analysed 10 days after the last administration. **(B)** Immunophenotypic analysis of *Padi4*^CTL^ and *Padi4*^CKO^ mice was performed 10 days after the final dose of 5-FU. Total number of cells in BM: WBC, LSK, and LK; HSC, MPP, HPC-1 and HPC-2 cells; myeloid, erythroid and lymphoid progenitor cells: CMP, GMP, MEP, CLP, Pre-GMP, Pre-MegE, MkP, Pre-CFU, CFU-E, Pro-Ery; differentiated Granulocytes, Monocytes, B cells, Erythroid cells. *Padi4*^CTL^, n = 21; *Padi4*^CKO^, n = 18. **(C, D)** Assessment of *Padi4* deletion on ageing of HSCs. **(C)** Immunophenotypic analysis in BM of *Padi4*^CTL^ and *Padi4*^CKO^ mice aged for 1 year. WBC, LSK, LK; HSC, MPP, HPC-1 and HPC-2 cells; myeloid, erythroid and lymphoid progenitor cells: CMP, GMP, MEP, CLP, Pre-GMP, Pre-MegE, MkP, Pre-CFU, CFU-E, Pro-Ery; differentiated Granulocytes, Monocytes, B cells, Erythroid cells. CFC assay with aged BM samples. Total number of cells in Spleen; WBC, Granulocytes, Monocytes, B cells, erythroid cells and T cells. *Padi4*^CTL^, n = 9; *Padi4*^CKO^, n = 8. **(D)** 200 CD45.2+ BM HSCs from 1 year old mice were transplanted to primary recipient mice and monitored for 24 weeks following which immunophenotypic analysis was performed on BM and spleen. Contribution of donor derived CD45.2+ cells to the Granulocyte, Monocyte, B cell and T cell population in PB. *Padi4*^CTL^, n = 20; *Padi4*^CKO^, n = 16. Percentage of donor-derived CD45.2+ cells in total BM, LSK, LK, HSC, MPP, HPC-1, HPC-1 and differentiated cell lineages (Granulocytes, Monocytes, B cells and erythroid cells). Contribution of donor-derived CD45.2+ cell population to total spleen WBC count and differentiated cell populations. *Padi4*^CTL^, n = 17; *Padi4*^CKO^, n = 15. 3-4 donors were used per genotype. All data are mean ± SEM. *, P < 0.05 (Mann-Whitney U test).

**Supplementary Figure 1. Blood profiling of *Padi4***^**CTL**^ **and *Padi4***^**CKO**^ **mice**. Automated cell counting of blood samples from 8-12 week old *Padi4*^CTL^ and *Padi4*^CKO^; WBC, RBC, HGB, HCT, MCV, MCH, MCHC and PLT counts. *Padi4*^CTL^, n = 8; *Padi4*^CKO^, n = 6. Data are mean ± SEM. *, P < 0.05; **, P < 0.01; ***, P < 0.001; ****, P < 0.0001 (Mann-Whitney U test).

**Supplementary Figure 2. Peripheral blood analysis of mice transplanted with *Padi4***^**CTL**^ **and *Padi4***^**CKO**^ **bone marrow**. Percentage of donor-derived CD45.2+ cells in PB and contribution of donor derived CD45.2+ cells to the Granulocyte, Monocyte, B cell and T cell population in PB. **(A)** Analysis of primary recipient mice. *Padi4*^CTL^, n = 36; *Padi4*^CKO^, n = 34. **(B)** Secondary recipient mice. *Padi4*^CTL^, n = 21; *Padi4*^CKO^, n = 22. Data are mean ± SEM. *, P < 0.05; **, P < 0.01; ***, P < 0.001; ****, P < 0.0001 (Mann-Whitney U test).

**Supplementary Figure 3. Acute deletion of *Padi4*. (A)** Deletion of *Padi4* following Poly I:C treatment in *Padi4*^IKO^ BM. **(B)** Percentage of donor-derived CD45.2+ cells in PB and contribution of donor derived CD45.2+ cells to the Granulocyte, Monocyte, B cell and T cell population in PB of recipient mice, after acute deletion of *Padi4*. n = 15–21 recipients per genotype. n = 3-4 donors per genotype. Data are mean ± SEM. *, P < 0.05; **, P < 0.01; ***, P < 0.001; ****, P < 0.0001 (Mann-Whitney U test).

**Supplementary Figure 4. 5-FU treatment leads to depletion of all bone marrow cells compartments in *Padi4***^**CTL**^ **mice**. Immunophenotypic analysis of *Padi4*^CTL^ mice treated with 5-FU or PBS vehicle control, demonstrating efficiency of 5-FU treatment. Experimental setup described in Figure 2A. Analysis was performed 10 days after the final dose of 5-FU; total number of cells in BM; WBC, LSK, and LK, HSC, MPP, HPC-1 and HPC-2 cells, myeloid, erythroid and lymphoid progenitor cells; CMP, GMP, MEP, CLP, Pre-GMP, Pre-MegE, MkP, Pre-CFU, CFU-E, Pro-Ery, differentiated Granulocytes, Monocytes, B cells, Erythroid cells and total number of cells in Spleen; WBC, B cells, Granulocytes and Monocytes *Padi4*^CTL^, n = 21; *Padi4*^CKO^ n = 18. Data are mean ± SEM. *, P < 0.05; **, P < 0.01; ***, P < 0.001; ****, P < 0.0001 (Mann-Whitney U test).

**Supplementary Figure 5. Peripheral blood analysis of mice transplanted with aged *Padi4***^**CTL**^ **and *Padi4***^**CKO**^ **bone marrow**. 200 CD45.2+ BM HSCs from 1 year old mice were transplanted to primary recipient mice and monitored for 24 weeks following which immunophenotypic analysis was performed on BM and spleen. Percentage of donor-derived CD45.2+ cells in PB. Contribution of donor derived CD45.2+ cells to the Granulocyte, Monocyte, B cell and T cell population in PB. *Padi4*^CTL^, n = 20; *Padi4*^CKO^, n = 16. Data are mean ± SEM. *, P < 0.05; **, P < 0.01; ***, P < 0.001; ****, P < 0.0001 (Mann-Whitney U test).

