## Supplementary Figures for "Peptidylarginine deiminase IV (PADI4) is not essential for cell-autonomous HSC maintenance and normal haematopoiesis"

#### Supplementary Figure 1

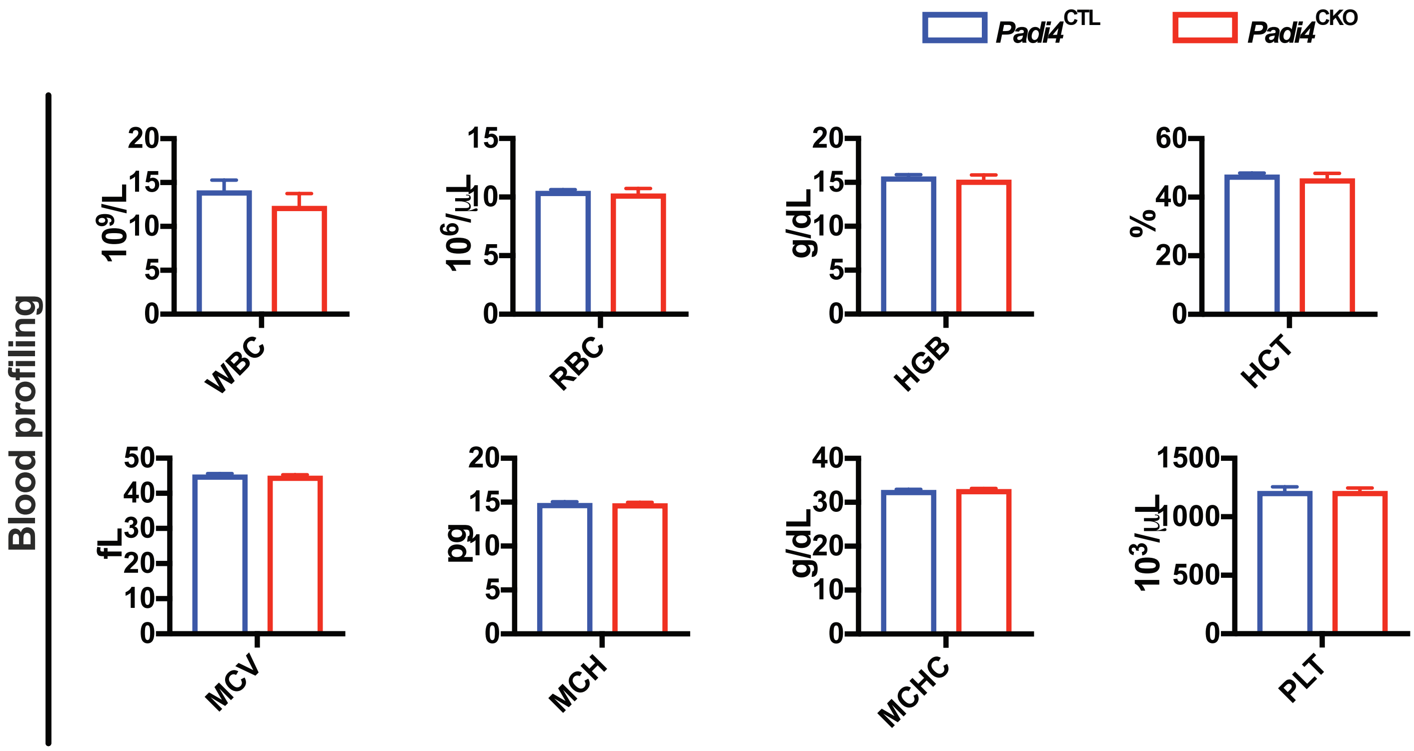

#### Supplementary Figure 2

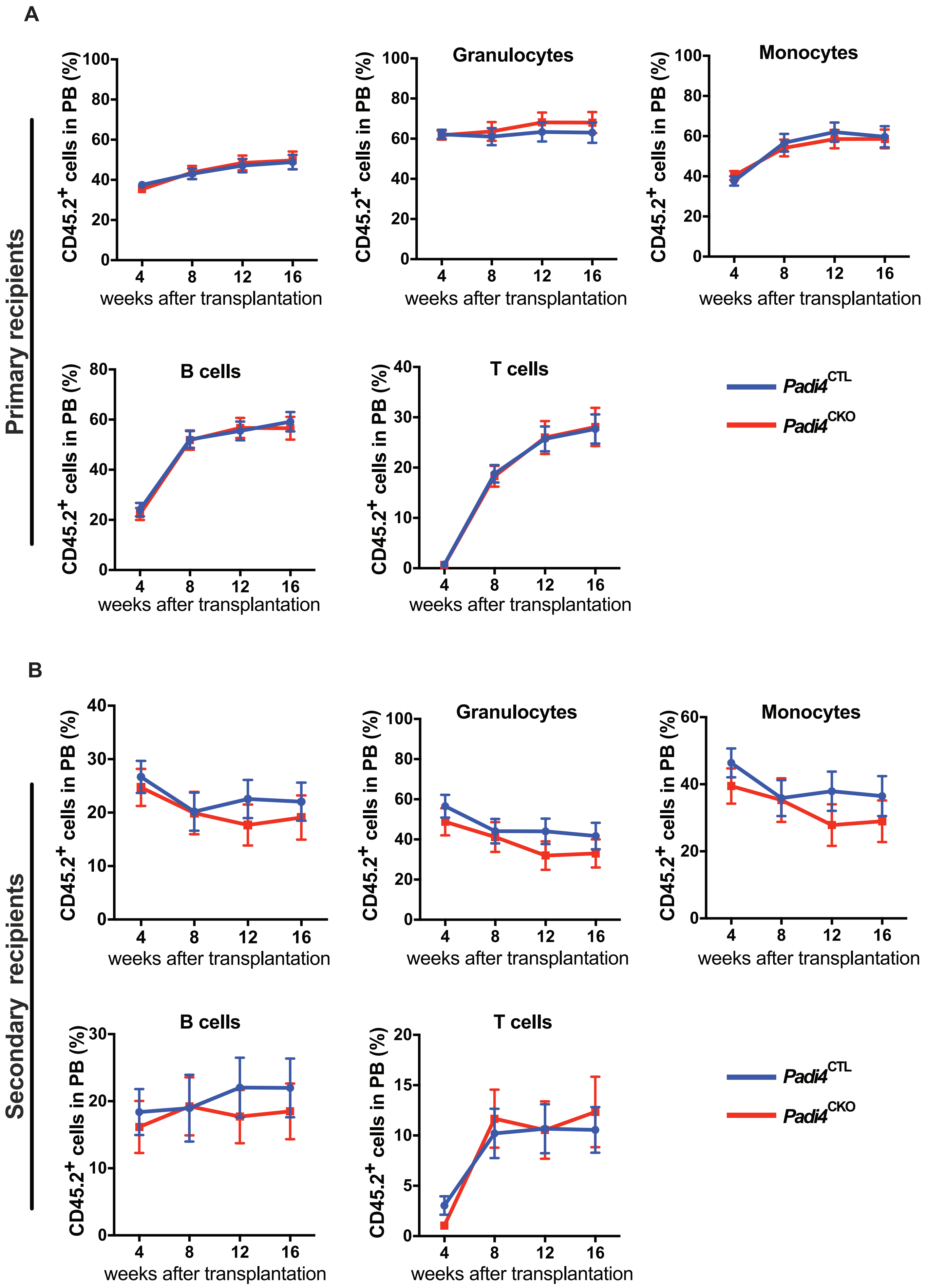

### Supplmenentary Figure 3

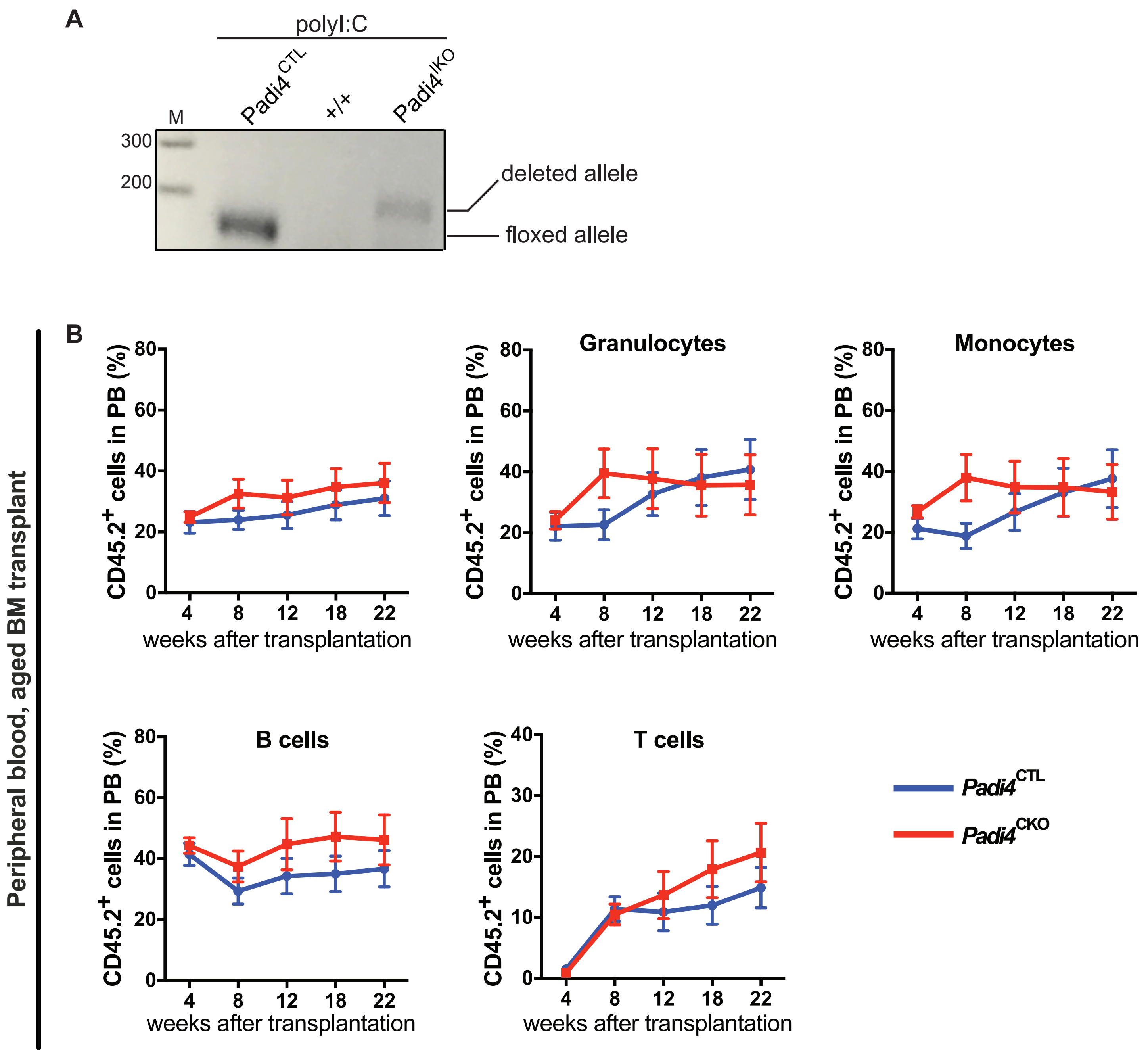

Supplmenentary Figure 4

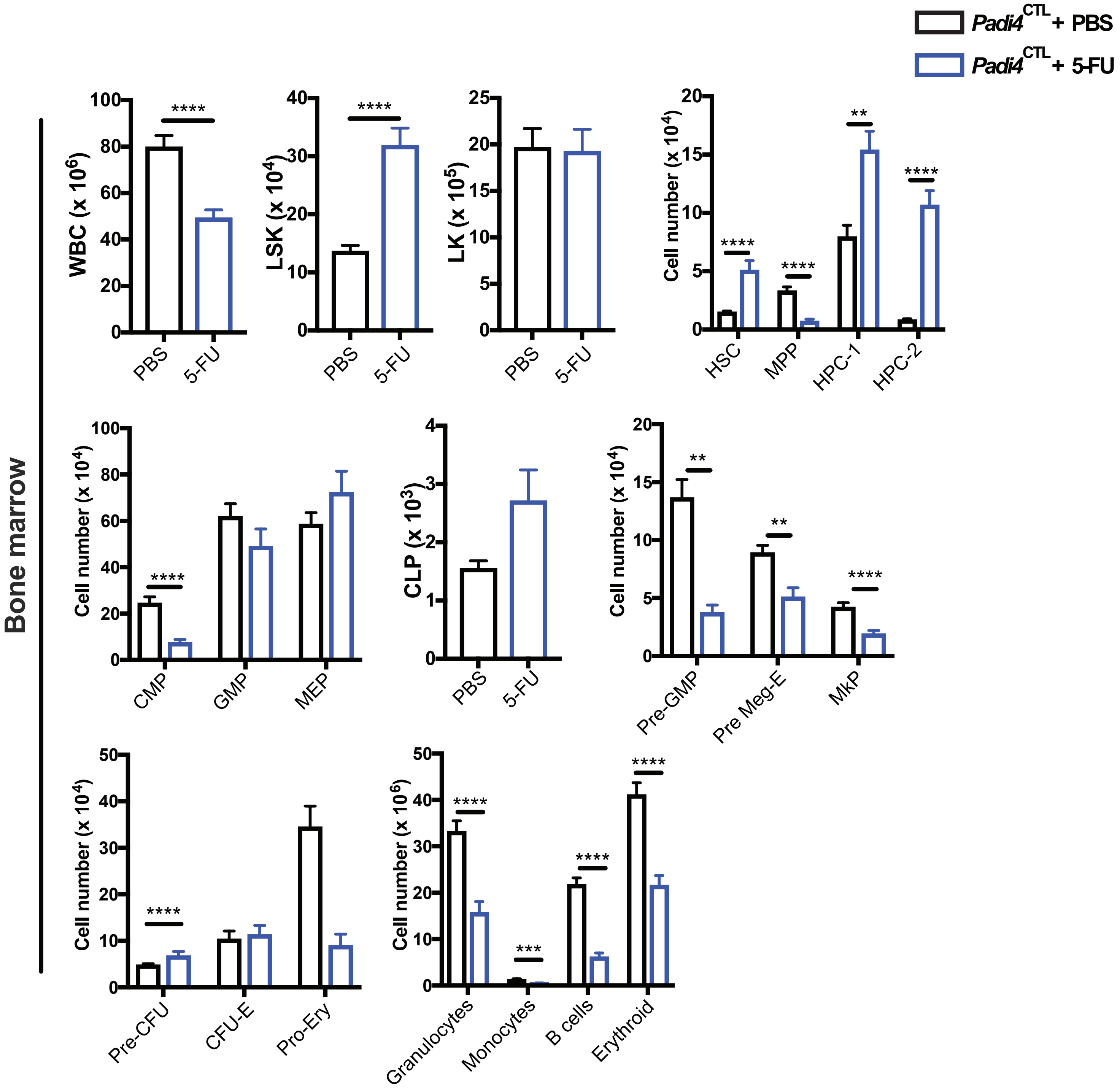

#### Supplmenentary Figure 5

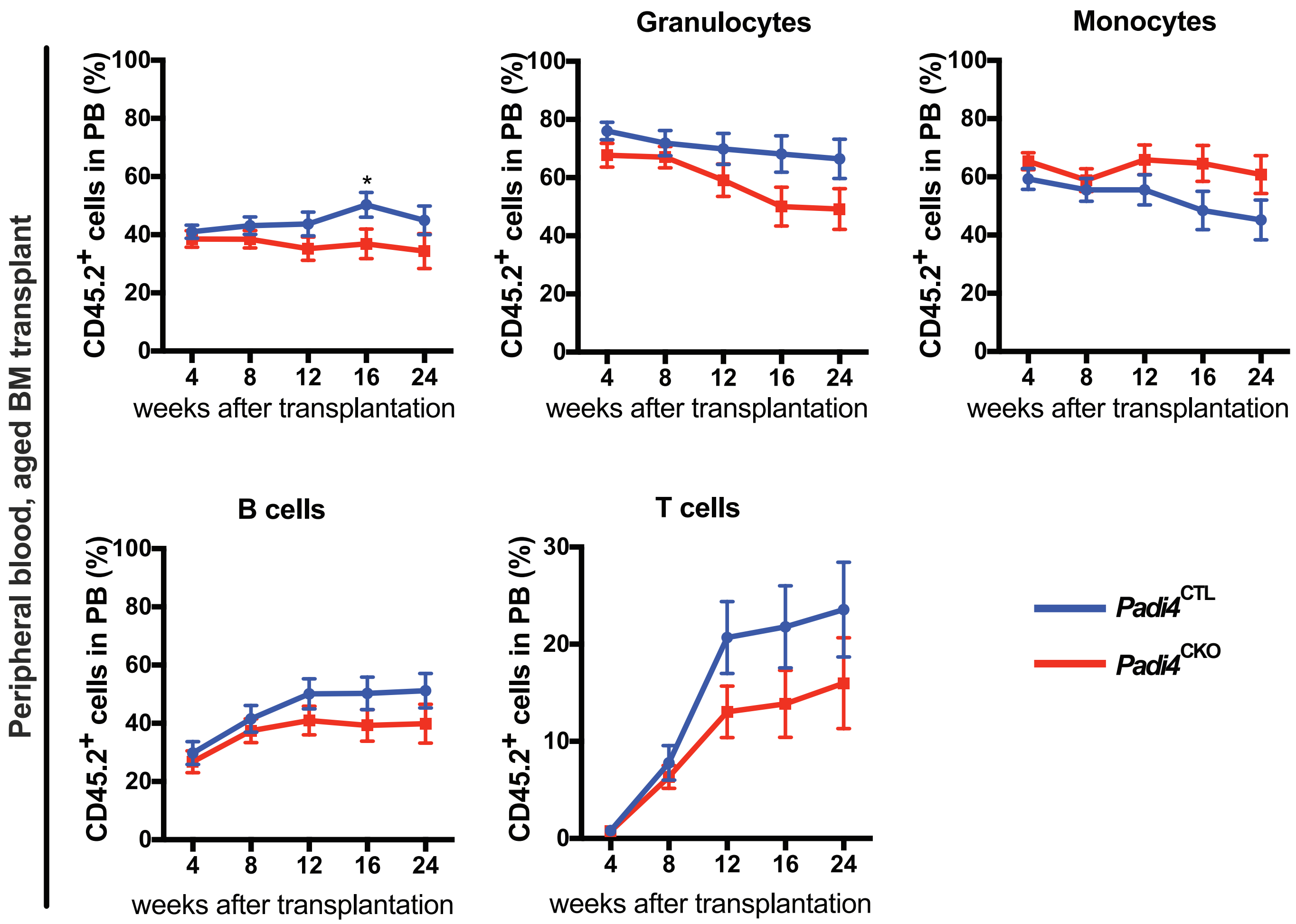
